## Supplementary Figures, Supplementary Tables and Supplementary Methods for "Rapid modulation of striatal cholinergic interneurons and dopamine release by satellite astrocytes"

#### Supplementary Figures S1-S14

#### Supplementary Tables S1-S3

#### Supplementary Methods – Mathematical Modelling

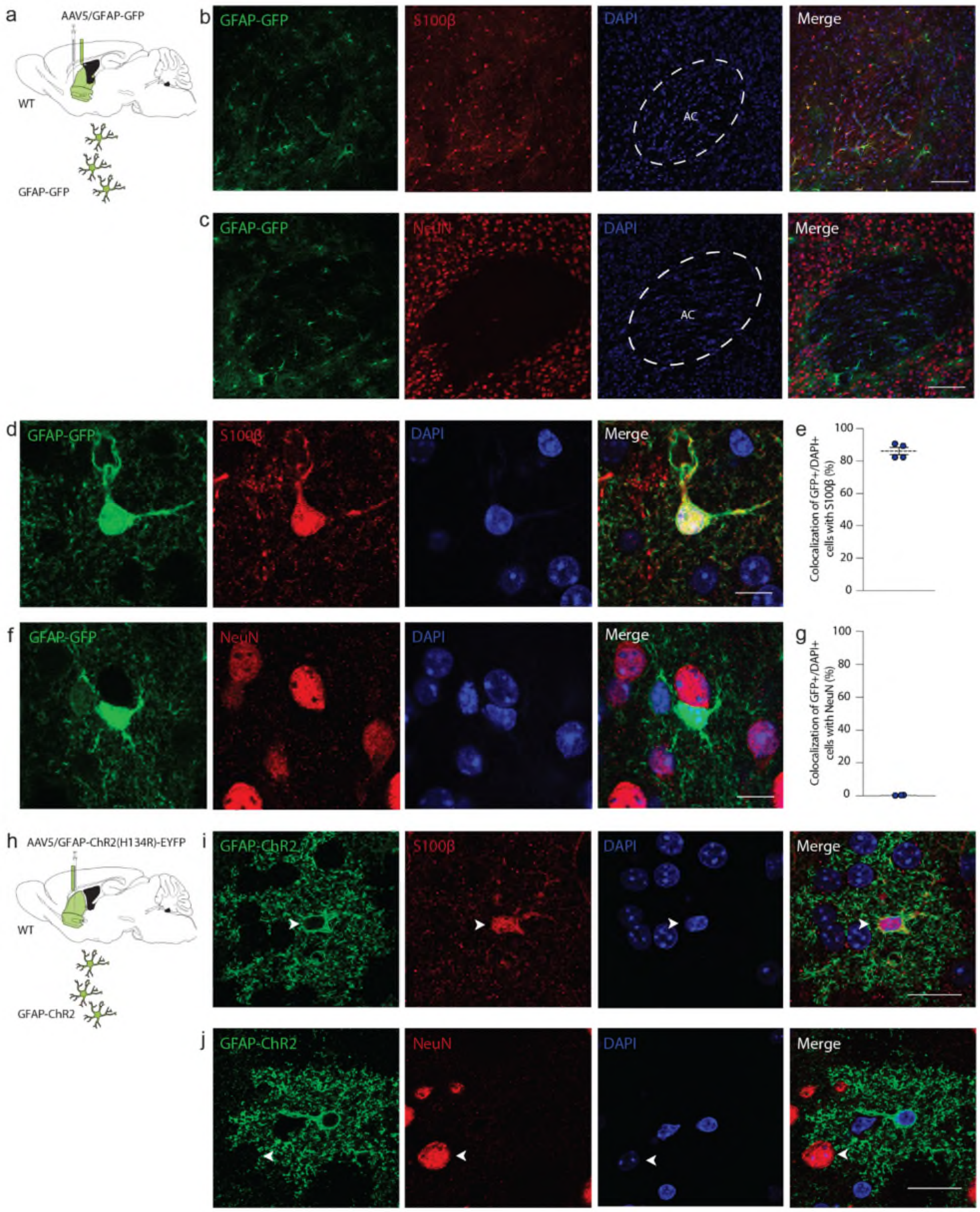


**SUPPLEMENTARY FIGURE S1. Specificity of gfabc1d ('GFAP') promotor driven viral transduction. a)** Experimental approach and location. WT mice received intracranial viral expression of GFAP-GFP in striatum, followed by immunofluorescent labeling and confocal microscopy ~3 weeks later. **b)** Low magnification single plane confocal image showing astrocytes co-labelled with GFAP-GFP (GFP; green) and S100β (red), along with DAPI+ to indicate nuclei (blue). Scale bar, 100 µm. **c)** Low magnification single plane confocal image showing astrocytes labelled with GFAP-GFP (GFP; green), neurons labelled with the neuronal marker NeuN (red), along with DAPI+ to indicate nuclei (blue). Scale bar, 100 µm. **d)** High magnification single plane confocal image showing an astrocyte labelled with GFAP-GFP (GFP; green) colocalizing with S100β (red), containing a DAPI+ nucleus (blue). Scale bar, 10 µm. **e)** Quantification of viral transduction specificity; the fraction of GFP+ cells that express the astrocyte marker S100β. *n* = 4 mice. **f)** High magnification single plane confocal image showing an astrocyte labelled with GFAP-GFP (GFP; green) not colocalizing with NeuN (red). Scale bar, 10 µm. **g)** Quantification of viral transduction specificity; the fraction of GFP+ cells that express the neuronal marker NeuN. *n* = 4 mice. **h)** Experimental approach and location. WT mice received intracranial viral expression of GFAP-ChR2-EYFP in striatum, followed by immunofluorescent labeling and confocal microscopy ~3-5 weeks later. **i)** High magnification single plane confocal image showing an astrocyte labelled with GFAP-ChR2-EYFP (EYFP; green) colocalizing with S100β (red; arrowhead), containing a DAPI+ nucleus (blue). Scale bar, 20 µm. **j)** High magnification single plane confocal image showing an astrocyte labelled with GFAP-ChR2-EYFP (EYFP; green) not colocalizing with NeuN (red; arrowhead). Scale bar, 20 µm. Error bars denote s.e.m. Abbreviations: AC, anterior commissure, WT, wildtype.

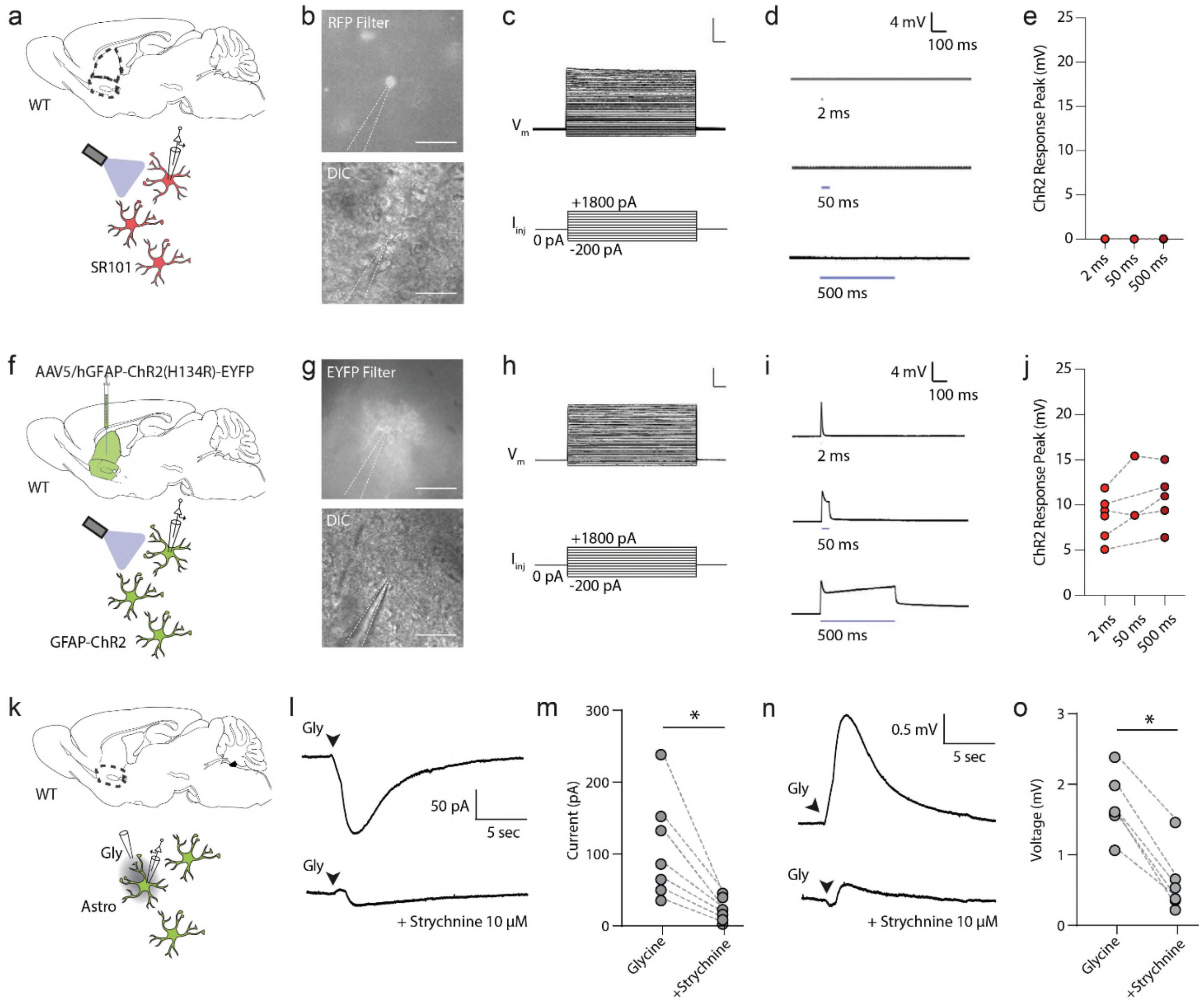


**SUPPLEMENTARY FIGURE S2. Optogenetic activation of ChR2-expressing but not wildtype astrocytes, and depolarization by striatal glycine. a)** Striatal wildtype astrocytes were labelled *ex vivo* throughout striatum with SR101, followed by whole-cell patch clamp electrophysiology. **b)** *Top*: Differential interference contrast (DIC) image showing whole-cell recording from a SR101-positive astrocyte. *Bottom*: RFP epifluorescent image showing the astrocyte positive for SR101. Scale bar, 50 µm. **c)** Astrocyte whole-cell current clamp recordings showing soma membrane voltage ($V_m$) responses under a 500-ms step current injections ($I_{inj}$), with a linear current-voltage relationship and absence of action potentials. **d-e)** Exposure to blue light (~4-5 mW, 2-50-500 ms) evokes no inward current/membrane voltage depolarization in SR101+ astrocytes ($n$ = 7(4)). **f)** Striatal astrocytes were virally labelled throughout striatum with GFAP-ChR2-EYFP, followed by *ex vivo* whole-cell patch clamp electrophysiology ~3-5 weeks later. **g)** *Top*: Differential interference contrast (DIC) image showing whole-cell recording from a ChR2-EYFP expressing astrocyte. *Bottom*: EYFP epifluorescent image showing the astrocyte positive for ChR2-EYFP. Scale bar, 50 µm. **h)** Astrocyte whole-cell current clamp recordings showing soma membrane voltage ($V_m$) responses under a 500-ms step current injections ($I_{inj}$), with a linear current-voltage relationship and absence of action potentials. **i-j)** Exposure to blue light (~4-5 mW, 2-50-500 ms) evokes an inward current/membrane voltage depolarization in EYFP+ astrocytes ($n$ = 6(3)). **k)** Whole-cell patch clamp electrophysiology of striatal wildtype astrocytes followed by local micropipette application of glycine (backfill 10 mM). **l-o)** Brief glycine puff elicited a potent and rapid inward current in voltage clamp (l-m) and depolarization by a few mV in current clamp (n-o), which was blocked by the classical glycine receptor antagonist strychnine (10 µM). * $p<0.05$ (Paired *t*-test). Abbreviations: Astro, astrocyte, $I_{inj}$ ,current injection, Gly, glycine, Vm, membrane voltage, WT, wildtype.

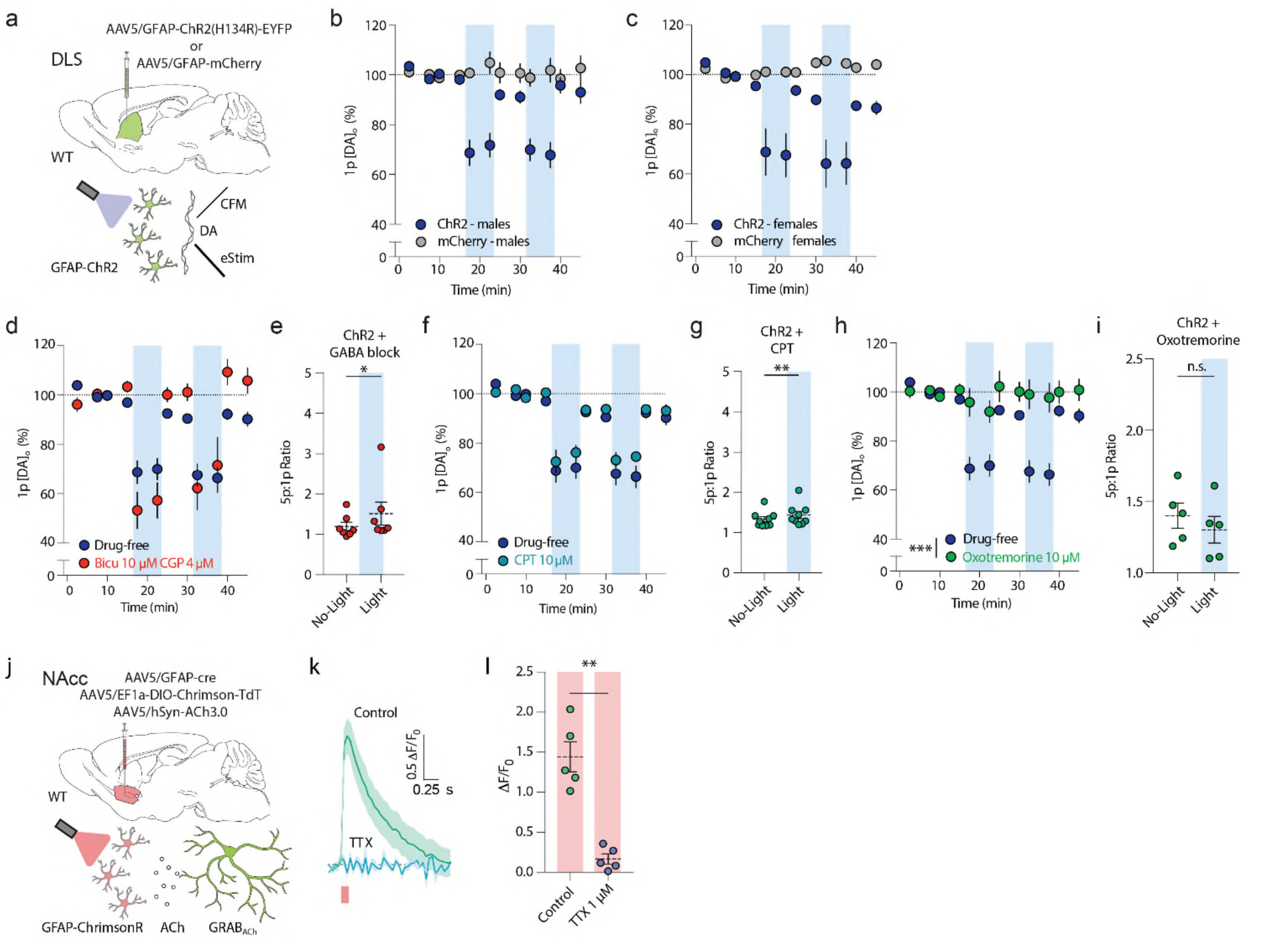


**SUPPLEMENTARY FIGURE S3. Similar dopamine release effects observed in male as well as female mice; absence of effects for GABA and adenosine antagonists. a)** Experimental overview. Striatal astrocytes were virally labelled in DLS with GFAP-ChR2-EYFP or GFAP-mCherry control followed by *ex vivo* fast-scan cyclic voltammetry combined with optogenetic activation (blue light; 500 ms; ~4-5 mW) ~3-5 weeks later. **a-c)** *Post hoc* segregation of the data between male (b) and female (b) mice show astrocyte-induced (blue column) alterations of dopamine release across both sexes. **d)** Summary of mean peak $[DA]_o$ evoked by a single electrical pulse (1p) before and under blue light (blue column) in GFAP-ChR2 mice (dark blue) normalized to pre-light baseline (dotted line), compared to combined application of $GABA_A$ receptor antagonist bicuculine (10 µM) and $GABA_B$ receptor antagonist CGP-55845 (4 µM; red). Application of GABA receptor antagonists does not prevent light induced reductions in dopamine release (Drug-Free: $n$ = 17(12), GABA block: $n$ = 7(5)). **e)** Ratio of peak $[DA]_o$ evoked by five consecutive pulses (5p; 50 Hz) compared to one electrical pulse (1p) at baseline and during 500-ms light activation (blue column) for ChR2 mice with application of $GABA_A$ receptor antagonist bicuculine (10 µM) and $GABA_B$ receptor antagonist CGP-55845 (4 µM). Blue light activation increased the 5p:1p ratio in ChR2 mice under blockade of GABA. GABA block: $n$ = 17(12). **f-g)** as (b)-(c) but with application of $A_1$ adenosine receptor antagonist CPT (10 µM). Blue light activation lowered DA release (f) and increased the 5p:1p ratio (g) in ChR2 mice under blockade of adenosine receptors. **h-i**) as (b)-(c) but with application of muscarinic M1 receptor agonist Oxotremorine (10 µM). Blue light activation (blue column) does not alter DA release (h) or the 5p:1p ratio (i) in ChR2 mice under blockade of muscarinic receptors, Oxo-M: $n = 8(5)$. **j-k)** Brief red-light activation of ChrimsonR-expressing astrocytes (10 ms; 4-5 mW, red column) evokes ACh release detected by $GRAB_{ACh3.0}$ sensor. **l)** Astrocyte-induced ACh release is TTX-sensitive (1 µM; $n$=5(5)). *** $p < 0.001$; ** $p < 0.01$; * $p < 0.05$. Repeated measures ANOVA in (d), (f), (h). Paired *t*-test in (e), (g), (i), (l). Abbreviations: bicu, bicuculline, CGP, CGP-55845 hydrochloride, ChI, cholinergic interneuron, CPT, 8-Cyclopentyl-1,3-dimethylxanthine, CFM, carbon fibre microelectrode, DA, dopamine, DLS, dorsolateral striatum, WT, wildtype.

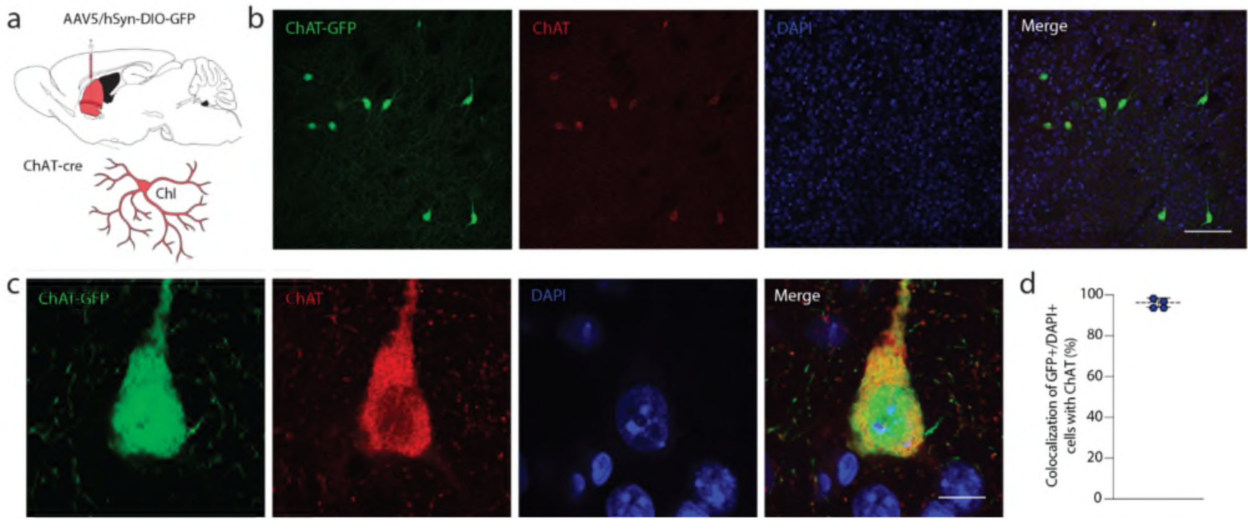


**SUPPLEMENTARY FIGURE S4. Validation of ChAT-cre transgenic line for viral transduction. a)** Experimental approach. *ChAT*-cre mice received intracranial viral expression of hSyn-DIO-mCherry or hSyn-DIO-eGFP throughout striatum, followed by histological examinations ~3-5 weeks later. **b)** Low magnification single plane confocal image showing an GFP-positive cell (ChAT-GFP; green) and antibodies against ChAT (red), along with DAPI+ to indicate nuclei (blue). Scale bar, 100 µm. **c)** High magnification single plane confocal image showing an eGFP-positive cell (ChAT-GFP; green) colocalizing with antibodies for ChAT (red), along with a DAPI+ nucleus (blue). Scale bar, 10 µm. **d)** Quantification of colocalization between GFP and ChAT antibodies. A large majority of GFP cells are positive for ChAT. *n* = 4. Error bars denote s.e.m.

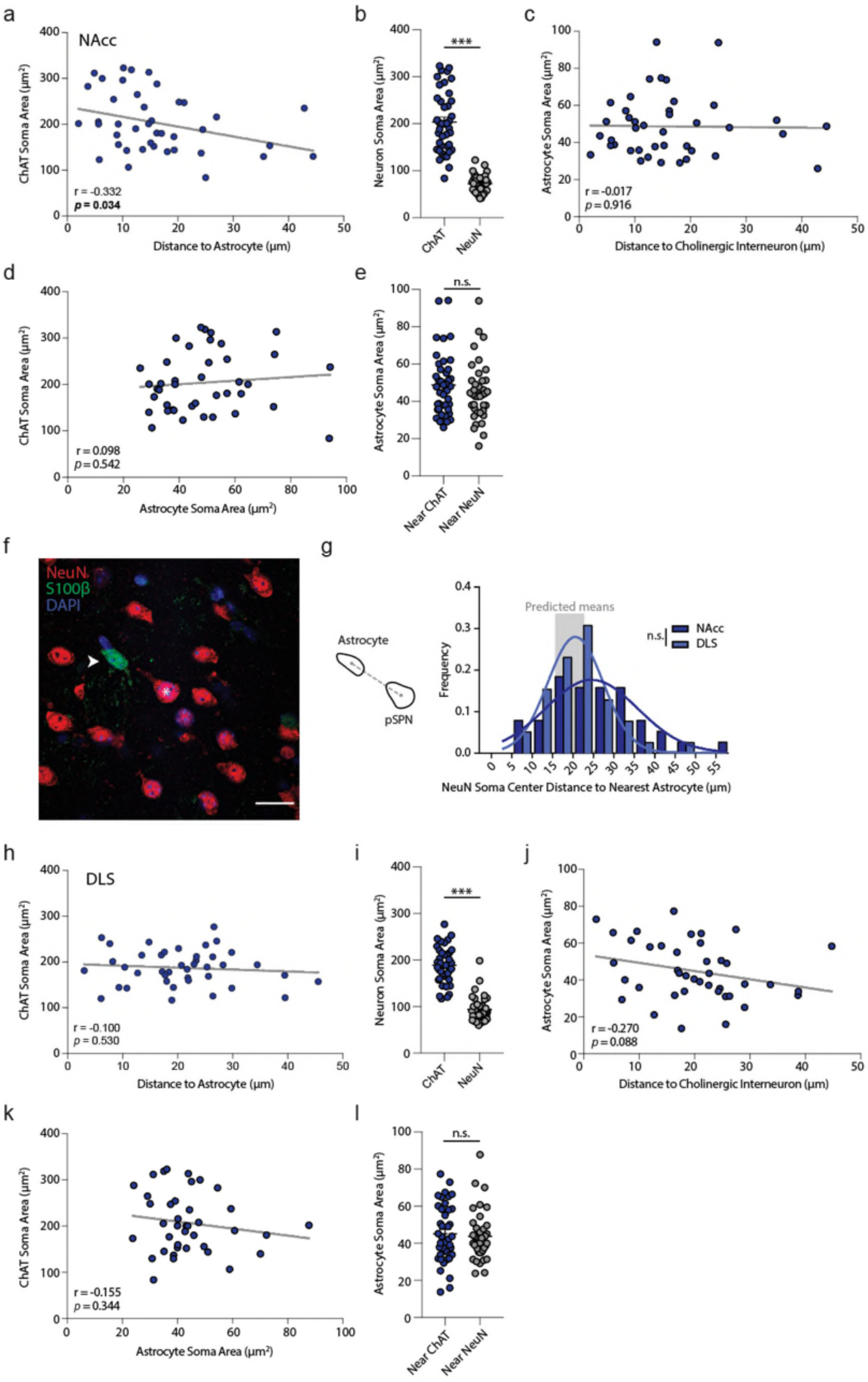


**SUPPLEMENTARY FIGURE S5. Correlations of astrocyte and cholinergic interneuron size and proximity; interaction between astrocytes and SPNs. a)** ChAT+ size correlates to distance of ChAT+ ChIs to nearest astrocytes in NAcc ($r$ = -0.332, $p$ = 0.034). **b)** Soma area of ChAT+ ChIs in mouse NAcc are larger than soma size of NeuN+ pSPNs, with minimal overlap ($t_{(77)}$ = 12, $p$ < 0.001, unpaired $t$-test). **c)** Astrocyte soma area in NAcc shows no correlation to their distance to ChI ($r$ = -0.017, $p$ = 0.916). **d)** ChAT+ soma size area in NAcc is not correlated to astrocyte soma area ($r$ = 0.098, $p$ = 0.542). **e)** Astrocyte soma area in NAcc does not differ per astrocytes found closest to either ChAT+ ChIs or NeuN+ pSPNs ($U$ = 675.5, $p$ = 0.313, Mann-Whitney $U$ test). **f)** Confocal microscopy image depicting astrocyte (GFAP, green) in proximity to a putative SPN (NeuN+, red). DAPI, blue. Scale bar, 20 µm. **g**) Left: Schematic depiction of the calculation of the soma-to-soma centre Eucidian distance. Right: Quantifications of observed astrocyte-pSPN soma-to-soma distances in DLS and NAc as compared to predicted distances. **h)** ChAT+ size does not correlate to distance of ChAT+ ChIs to nearest astrocytes in DLS ($r$ = -0.100, $p$ = 0.530). **i)** Soma area of ChAT+ ChIs in DLS are larger than soma size of NeuN+ pSPNs, with minimal overlap ($U$ = 43, $p$ < 0.001, Mann-Whitney $U$ test). **j)** Astrocyte soma area in DLS shows no correlation to their distance to ChI ($r$ = -0.270, $p$ = 0.088). **k)** ChAT+ soma size area in DLS is not correlated to astrocyte soma area ($r$ = -0.155, $p$ = 0.344). **l)** Astrocyte soma area in DLS does not differ per cells adjacent to ChAT+ ChIs or NeuN+ pSPNs ($U$ = 733.5, $p$ = 0.652, Mann-Whitney $U$ test). *** $p$ < 0.001; n.s. non-significant. Error bars denote s.e.m. Unpaired $t$-test in (b). Mann-Whitney $U$ test in (e), (i), (l). Pearson correlation in (a), (c), (d), (h), (j), (k). Abbreviations: DLS dorsolateral striatum, NAc nucleus accumbens core, pSPN putative spiny projection neuron.

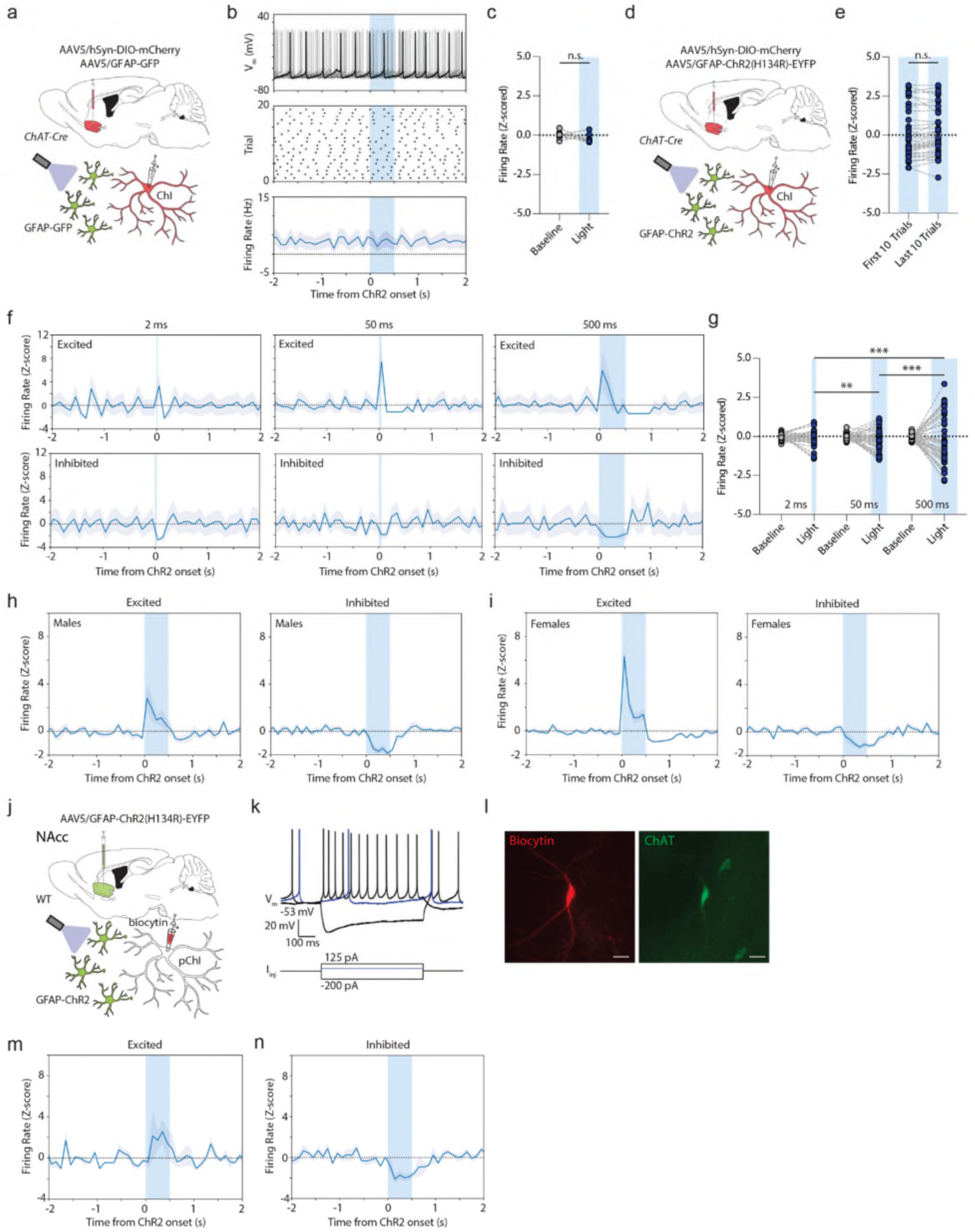


**SUPPLEMENTARY FIGURE S6. Absence of light only effects and presence of light duration-dependent modulation of cholinergic interneuron firing; bidirectional astrocyte activation of ChIs in wildtype mice. a)**

Experimental approach. *ChAT*-cre mice received intracranial viral expression of GFAP-GFP and hSyn-DIO-mCherry in NAcc, followed by whole-cell patch clamp electrophysiology and blue light activation ~3-5 weeks later. **b**) Raw membrane voltage trace (*top*), action potential scatter plot (*middle*), and spontaneous firing frequency (100 ms bins; *bottom*) for 20 consecutive trials for various example ChI, plotted from light onset. In response to blue light activation (~4-5 mW; 500 ms; blue column), ChI do not change their spontaneous firing. **c**) Z-scored firing rates for ChI during baseline (1000 ms before light onset) and during light activation (500-ms; $n$ = 10(4)). Blue light does not result in alterations in spontaneous firing frequency. **d)** Experimental approach. *ChAT*-cre mice received intracranial viral expression of hGFAP-ChR2-EYFP and hSyn-DIO-mCherry in NAcc, followed by whole-cell patch clamp electrophysiology and brief astrocyte optogenetic activation ~3-5 weeks later. **e)** Z-scored firing rate responses in ChIs in response to astrocyte optogenetic activation (500 ms; blue columns), segregated between the first 10 and last 10 trials. Response direction (positive or negative) is stable over 20 consecutive trials. **f)** Z-scored firing rate responses in ChIs grouped to their response direction to astrocyte optogenetic activation (blue column), which show deflections from baseline firing rate for 2 ms light activation (*left*), 50 ms light activation (middle), and 500 ms light activation for excitatory responses (*top*) and inhibitory responses (*bottom*). **g)** Z-scored firing rates for ChIs during baseline (1000 ms before light onset) and during light activation (blue colum, $n$ = 41(13)) for varying light durations. $p$ = 0.001, 2 ms versus 50 ms; $p$ < 0.001, 50 ms versus 500 ms; $p$ < 0.001, 2 ms versus 500 ms; One-way ANOVA followed by Tukey's multiple comparisons. **h-i)** Z-scored firing rate responses in ChIs in response to light activation (500 ms, blue column) segregated by their response direction, as well as *post hoc* segregated for male (h) and female (i) mice. Male excitation $n$ = 6(5), inhibition $n$ = 12(6); female excitation $n$ = 8(5) female inhibition $n$ = 9(5). **j)** Experimental approach. Wildtype (WT) mice received intracranial viral expression of GFAP-ChR2-EYFP in NAcc, followed by whole-cell patch clamp electrophysiology (and biocytin filling) of putative ChIs and astrocyte optogenetic activation ~3-5 weeks later. **k)** Patch clamp electrophysiological recordings of putative ChIs in current clamp configuration (*top* trace) in response to square negative (black), positive (black), and threshold (blue) current injections. Putative ChIs show rhythmic spontaneous activity, depolarized membrane potential, voltage sag in response to negative current injections, and a large afterhyperpolarization, similar to Fig. 3c. Scale bar, 20 mV. **l)** Single plane confocal microscopy image showing biocytin-filled ChI (red) and *post hoc* immunofluorescent labeling for ChAT (green). Scale bar, 20 µm. **m-n)** Z-scored firing rate responses in ChIs in response to blue light activation (500 ms, blue column) grouped to their response direction, segregated for excitation responses (m) and inhibition responses (n) mice. Shaded areas denote s.e.m. *** $p$ < 0.001; ** $p$ < 0.01. Paired $t$-test in (c), (e). One-way ANOVA followed by Tukey's multiple comparisons in (g). Error bars and shaded areas denote s.e.m. Abbreviations: Astro astrocyte, ChI cholinergic interneuron, n.s. non-significant, pChI putative cholinergic interneuron, $V_m$ membrane voltage, WT wildtype.

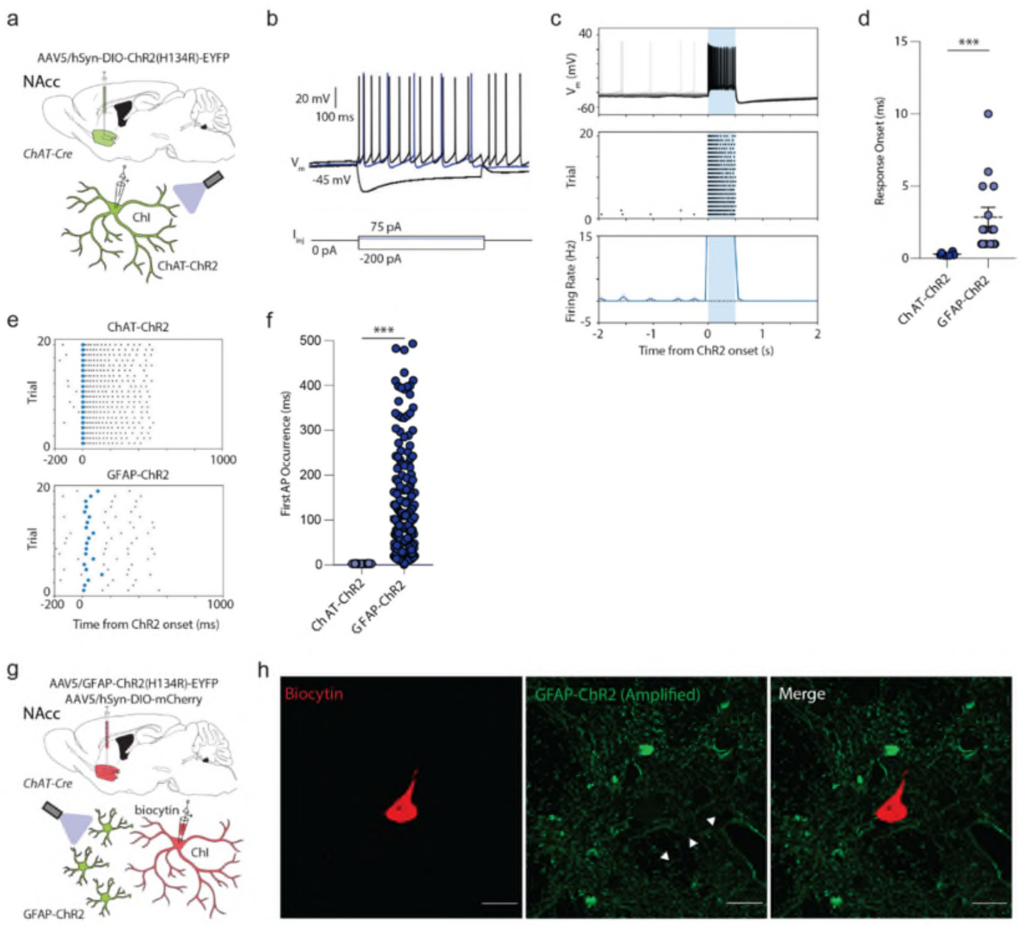


**SUPPLEMENTARY FIGURE S7. No evidence for presence of ChR2-EYFP in ChIs. a)** Experimental overview. **b)** Patch-clamp recording of targeted ChI, current clamp configuration (*top* trace) in response to square negative (black), positive (black), and threshold (blue) current injections. ChIs show rhythmic spontaneous activity, depolarized membrane potential, voltage sag in response to negative current injections, and a large afterhyperpolarization. **c)** Raw membrane voltage trace (*top*), action potential scatter plot (middle), and firing frequency (100 ms bins; bottom) for 20 consecutive trials for ChIs, plotted from light onset (~4-5 mW; 500 ms). **d)** Quantification of response onsets between blue light-induced responses in ChIs with ChR2 expressed in astrocytes (GFAP-ChR2; corresponding to Fig. 1j) or in ChIs themselves (ChAT-ChR2). ($U$ = 0.000, $p$ < 0.001, Mann-Whitney $U$ test). **e)** Scatter plot, action potentials in ChIs in response to 500-ms blue light activation in representative cells from ChAT-ChR2 mice (*top*) versus GFAP-ChR2 mice (bottom) over 20 trials. First elicited action potential following light onset is indicated in blue. Note the larger intertrial variability for elicited action potentials within a single cell in GFAP-ChR2 mice. **f)** Quantification of first action potential onset variation in ChAT-ChR2 versus GFAP-ChR2 cells ($U$ = 488, $p$ < 0.001, Mann-Whitney $U$ test). **g)** Experimental overview. **h)** High magnification maximum projection of biocytin-filled ChIs together with GFP antibody-enhanced GFAP-ChR2-EYFP, showing close colocalization but no overlap (arrowheads) between the ChR2 and labelled ChI. Scale bar, 20 µm. *** $p$ < 0.001, ** $p$ < 0.01. Mann-Whitney $U$ test in (d) and (f).

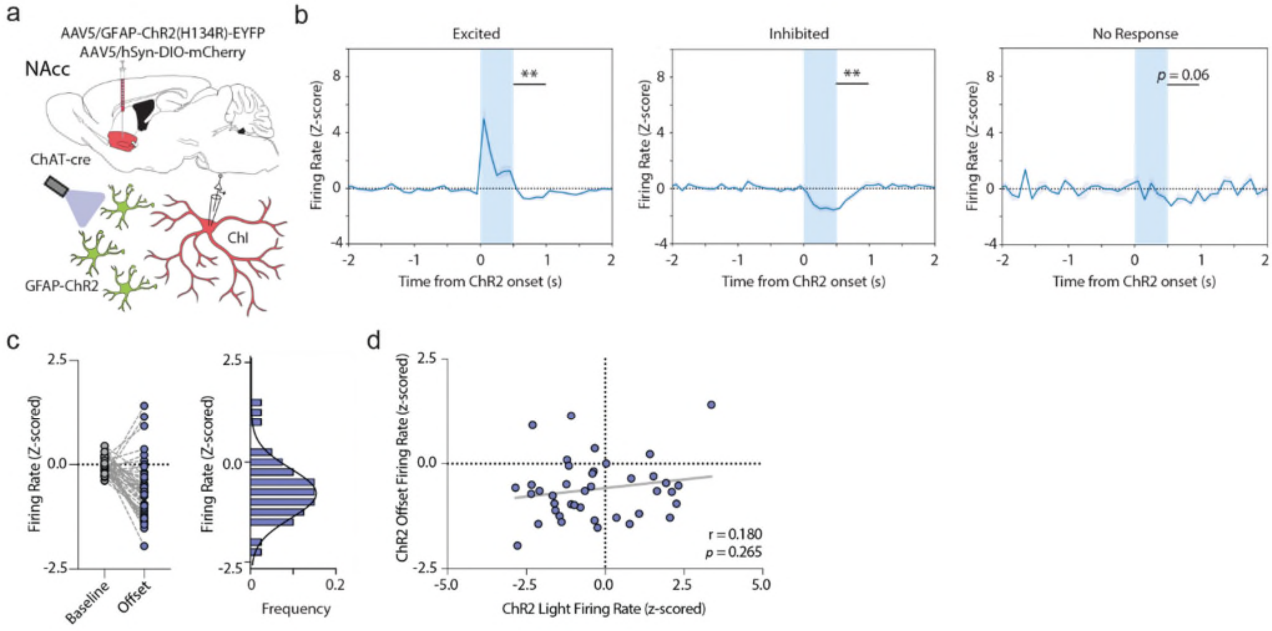


**SUPPLEMENTARY FIGURE S8. Unimodal inhibitory responses across ChIs following astrocyte activation a)** Experimental approach. *ChAT*-cre mice received intracranial viral expression of hGFAP-ChR2-EYFP and hSyn-DIO-mCherry in NAcc, with astrocyte optogenetic activation during targeted whole-cell patch clamp recordings of ChIs in NAcc ~3-5 weeks later. **b)** Corresponding to Fig. 3. Z-scored spontaneous firing rates for ChIs during light activation ($n$ = 41(13); blue bar) for excited cells (*left*), inhibited cells (*middle*), and no response cells (*right*). Z-scored firing rate responses after light offset of ChIs show reductions in firing rate for ~500 ms (*left*: $t_{(11)} = 3.866$, $p = 0.003$; paired *t*-test; *mid*: $t_{(20)} = 3.841$, $p = 0.001$; paired *t*-test; *right*: $t_{(7)} = 2.250$, $p = 0.060$; paired *t*-test). **c)** *Left*: Z-scored firing rates for ChIs during baseline (1000 ms before light onset) and after light offset (500 ms) ($n$ = 41(13)). *Right*: Z-scored firing rate responses after light offset of ChIs show unimodal reductions in firing rate for ~500 ms. **d)** Scatter plot of ChI z-scored firing rates during blue light onset (500 ms) and during a 500 ms window following light offset, fitted with a linear function (grey line). No correlation of z-scored firing rates during light and following light offset were observed ($r = 0.186$, $p = 0.265$). ** $p < 0.01$. Paired *t*-test in (b). Pearson correlation in (d). Error bars and shaded areas denote s.e.m. Abbreviations: AP action potential, ChI cholinergic interneuron, NAcc nucleus accumbens core, n.s. non-significant, $I_{inj}$ current injection, $V_m$ membrane potential.

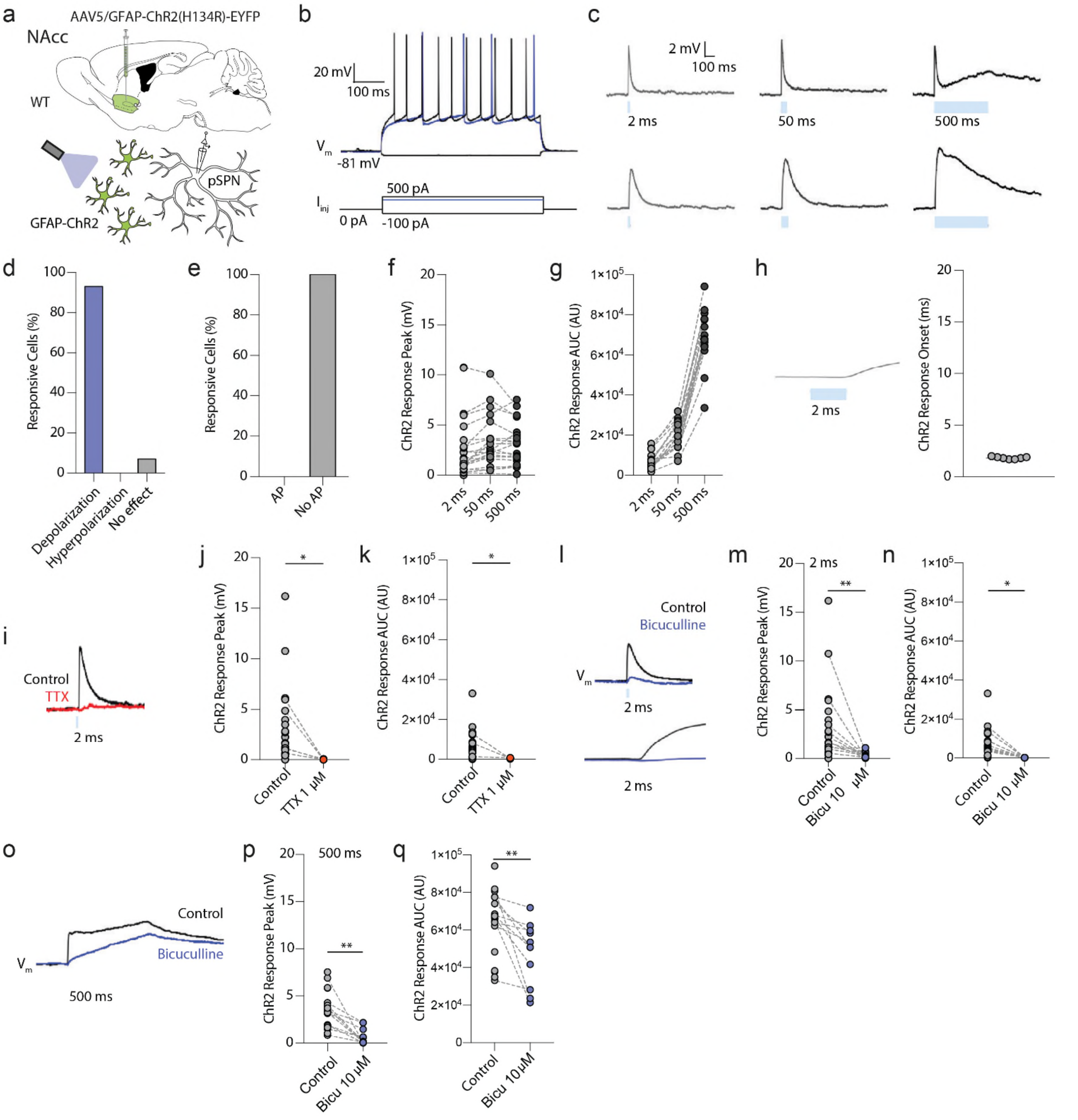


**SUPPLEMENTARY FIGURE S9. Astrocyte activation induces small unimodal TTX-sensitive depolarization responses in pSPNs; astrocyte effects on pSPNs are prevented by GABA$_A$ receptor antagonists. a)** Experimental approach. Striatal astrocytes were virally labelled in NAcc with GFAP-ChR2-EYFP, followed by *ex vivo* whole-cell patch clamp electrophysiology of putative SPNs ~3-5 weeks later. **b)** Patch-clamp electrophysiological recordings of targeted pSPNs in NAcc in current clamp configuration (*top* trace) in response to square negative (black), positive (black), and threshold (blue) current injections. No voltage sag in response to negative current injections, low input resistance, delayed response to first action potential and broad action potentials. Scale bars, 20 mV, 100 ms. **c)** Membrane voltage traces for two pSPNs in response to various durations of astrocyte activation (4-5 mW, blue light). Scale bars, 2 mV, 100 ms. **d)** Quantification of response direction of pSPNs after astrocyte activation. **e)** Quantification of elicited action potentials. No cells showed any *de novo* action potentials in response to astrocyte activation. **f)** Quantification of membrane voltage peak depolarization in response to astrocyte activation (2-50-500 ms). **g)** Quantification of membrane voltage AUC depolarization in response to astrocyte activation of different durations (2-50-500 ms). **h)** *Left*: Example membrane voltage deflection of a pSPN in response to astrocyte activation (blue bar). *Right*: quantification of response onset in response to 2 ms astrocyte activation. **i)** Membrane voltage traces in response to 2 ms of astrocyte activation before (black) and after application of TTX (1 µM; red). Scale bar, 4 mV, 2 ms. **j)** Quantification of peak membrane voltage responses with application of TTX (1 µM). Application of TTX prevents the responses in pSPNs induced by 2 ms astrocyte activation. **k)** Quantification of voltage response AUC with application of TTX (1 µM). TTX prevents the response in pSPNs. **l)** Membrane voltage traces in response to 2 ms of astrocyte activation before (black) and after application of the specific $GABA_A$ receptor antagonist bicuculline (10 µM; blue). Scale bar, 2 mV, 2 ms. **m)** Quantification of peak membrane voltage responses with application of bicuculline (10 µM). Independent application of bicuculline prevents the responses in pSPNs induced by 2 ms astrocyte activation. **n)** Quantification of voltage response AUC with application of bicuculline (10 µM). Application of bicuculline prevents the response AUC in pSPNs induced by 2 ms astrocyte activation. **o)** Membrane voltage traces in response to 500 ms of astrocyte activation before (black) and after application of the specific $GABA_A$ receptor antagonist bicuculline (10 µM; blue). Scale bar, *2* mV, 2 ms. **p)** Quantification of peak membrane voltage responses with application of bicuculline (10 µM). Application of bicuculline prevents the initial peak response in pSPNs induced by 500 ms astrocyte activation. **q)** Quantification of voltage response AUC with application of bicuculline (10 µM). Application of bicuculline lowers but does not fully prevent the response AUC in pSPNs induced by 500 ms astrocyte activation. *** $p < 0.001$, ** $p < 0.01$, * $p < 0.05$. Paired t-test in (j), (k), (o), (q). Error bars denote s.e.m. Abbreviations: AP action potential, AUC area under the curve, ChI cholinergic interneuron, $I_{inj}$ current injection, NAcc nucleus accumbens core region, pSPN putative spiny projection neuron, $V_m$ membrane voltage, WT wildtype.

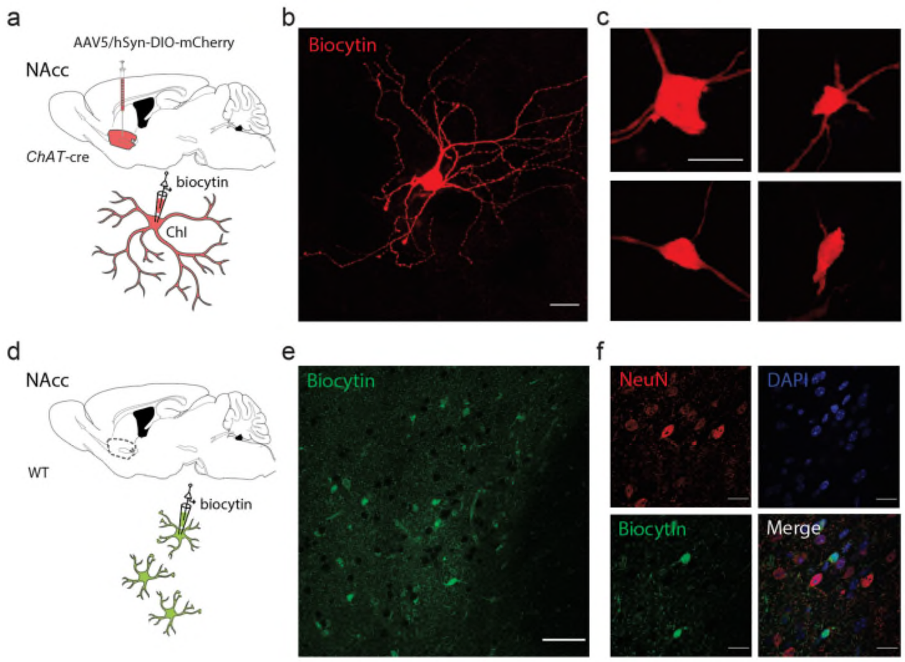


**SUPPLEMENTARY FIGURE S10. Astrocyte excitation of ChIs is synapse- and gap junction-independent. a)** Experimental overview. *ChAT*-cre mice received intracranial viral expression of hSyn-DIO-mCherry in NAcc, followed by targeted whole-cell patch clamp recordings and biocytin filling of ChIs ~3-5 weeks later. **b)** Maximum projection of a confocal microscopy image of a biocytin-filled ChI (red). Biocytin filling of ChIs does not result in labelling of astrocytes. Scale bar, 10 µm. **c)** Additional examples of biocytin-filled ChIs without notable labeled glial cells. Scale bar, 20 µm. **d)** Experimental overview. Targeted whole-cell patch clamp recordings guided by the glial dye SR101 and consequent biocytin filling of astrocytes in NAcc. **e)** Biocytin filling of single astrocytes leads to widespread labelling of adjacent astrocytes. Scale bar, 40 µm. **f)** Single plane colocalization images showing biocytin-labeled astrocytes (green), along with labeled neurons (NeuN; red), and DAPI+ nuclei (blue). Scale bar, 20 µm. No biocytin-labeled neurons were observed.

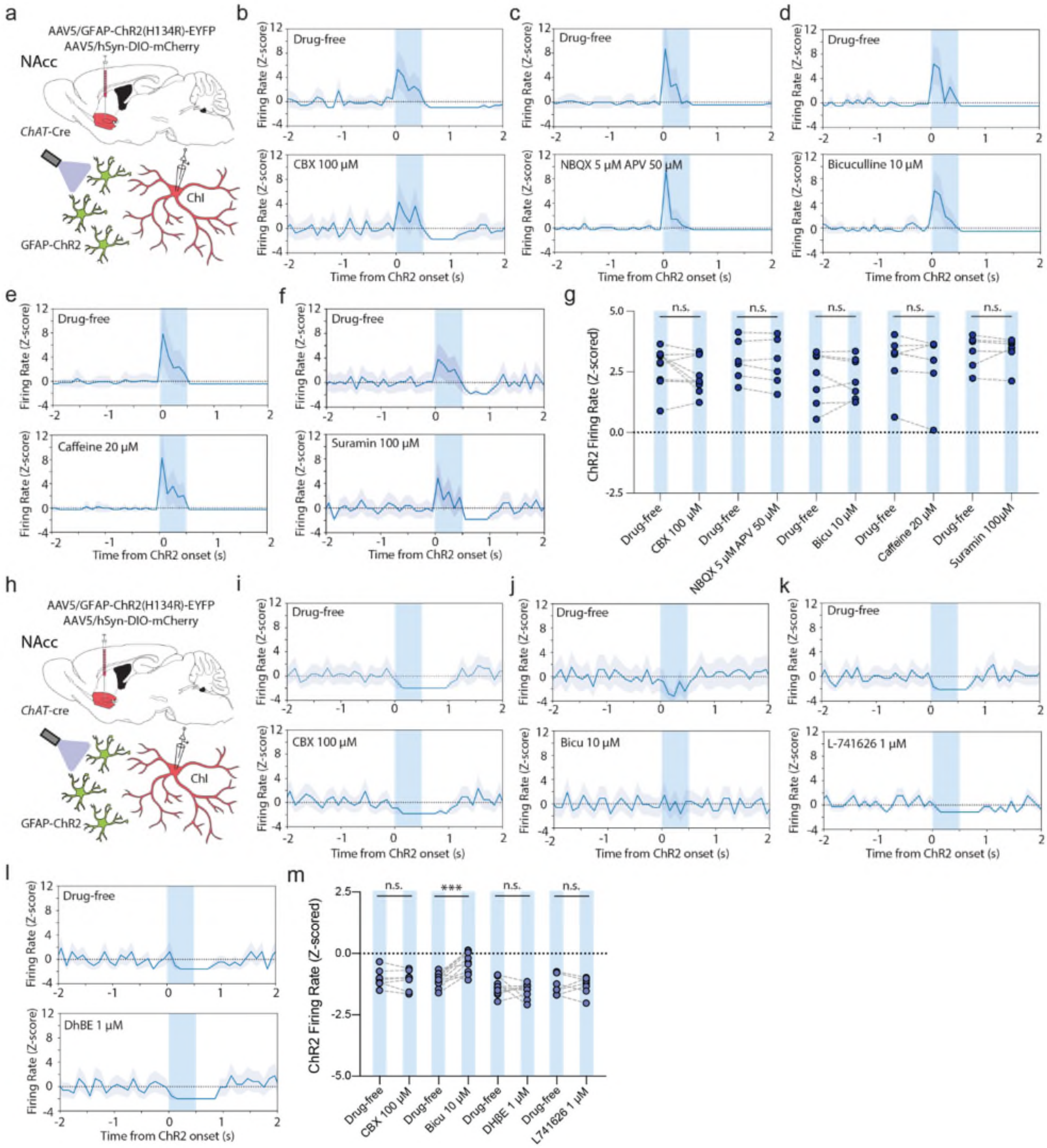


**SUPPLEMENTARY FIGURE S11. Astrocytes inhibit ChIs through feedforward GABAergic inhibition. a)** Experimental approach. **b-f)** Z-scored firing rate responses in an example excitation ChI under 500-ms blue light activation (blue column) in drug-free conditions (*top*) and drug application (bottom), for application of the non-selective gap junction blocker carbenoxolone (CBX, 100 µM, b), application of the NMDA and AMPA receptor antagonists, NBQX (5 µM) and AP-V (50 µM, c), application of $GABA_A$ receptor antagonist bicuculline (10 µM, d), application of adenosine $A_1$ and $A_2$ receptor antagonist caffeine (20 µM, e), and application of purinergic receptor antagonist suramin (100 µM, f). **g)** Z-scored excitation firing rate changes for drugs under light activation in drug-free and drug application show no effect for applied drugs. Carbenoxolone (*n* = 9(6)). NBQX and APV (*n* = 6(3)). bicuculline (*n* = 7(5)). caffeine (*n* = 6(3)). Suramin (100 µM). (*n* = 6(3)). **h)** Experimental approach. **i-l)** Z-scored firing rate responses in an example excitation ChI under 500-ms blue light activation (blue column) in drug-free conditions (*top*) and drug application (bottom), for application of the non-selective gap junction blocker carbenoxolone (CBX, 100 µM, i), application of the $GABA_A$ receptor receptor antagonist bicuculline (10 µM, j), application of dopamine D2 receptor receptor antagonist L-741626 (L-741626, 1 µM, k), or application of nAChR receptor antagonist DHβE (1 µM, l). **m)** Z-scored excitation firing rate changes for drugs under light activation in drug-free and drug application are prevented by GABAa block. Carbenoxolone (*n* = 8(6)). Bicuculline (*n* = 11(7)). L-741626 (*n* = 7(3)). DHβE (*n* = 9(5)). *** $p < 0.001$; Paired *t*-tests in (g), (m). Shaded areas denote s.e.m. Abbreviations: Astro astrocyte, bicu bicuculline, CBX carbenoxolone, ChI cholinergic interneuron, Exci excitation, NAcc nucleus accumbens core region, n.s. non-significant, WT wildtype.

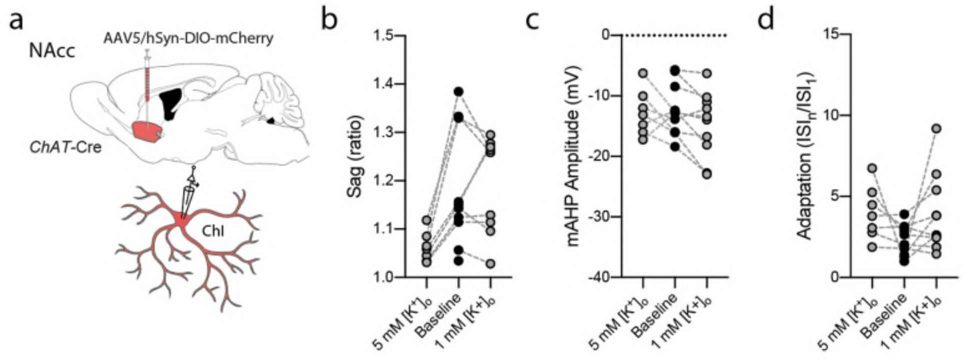


**SUPPLEMENTARY FIGURE S12. a-d)** Changing extracellular aCSF $[K^+]_o$ from baseline (1.5 mM) to 1 mM or 5 mM has non-linear effects on the sag ratio, mAHP amplitude, and spike frequency adaptation.

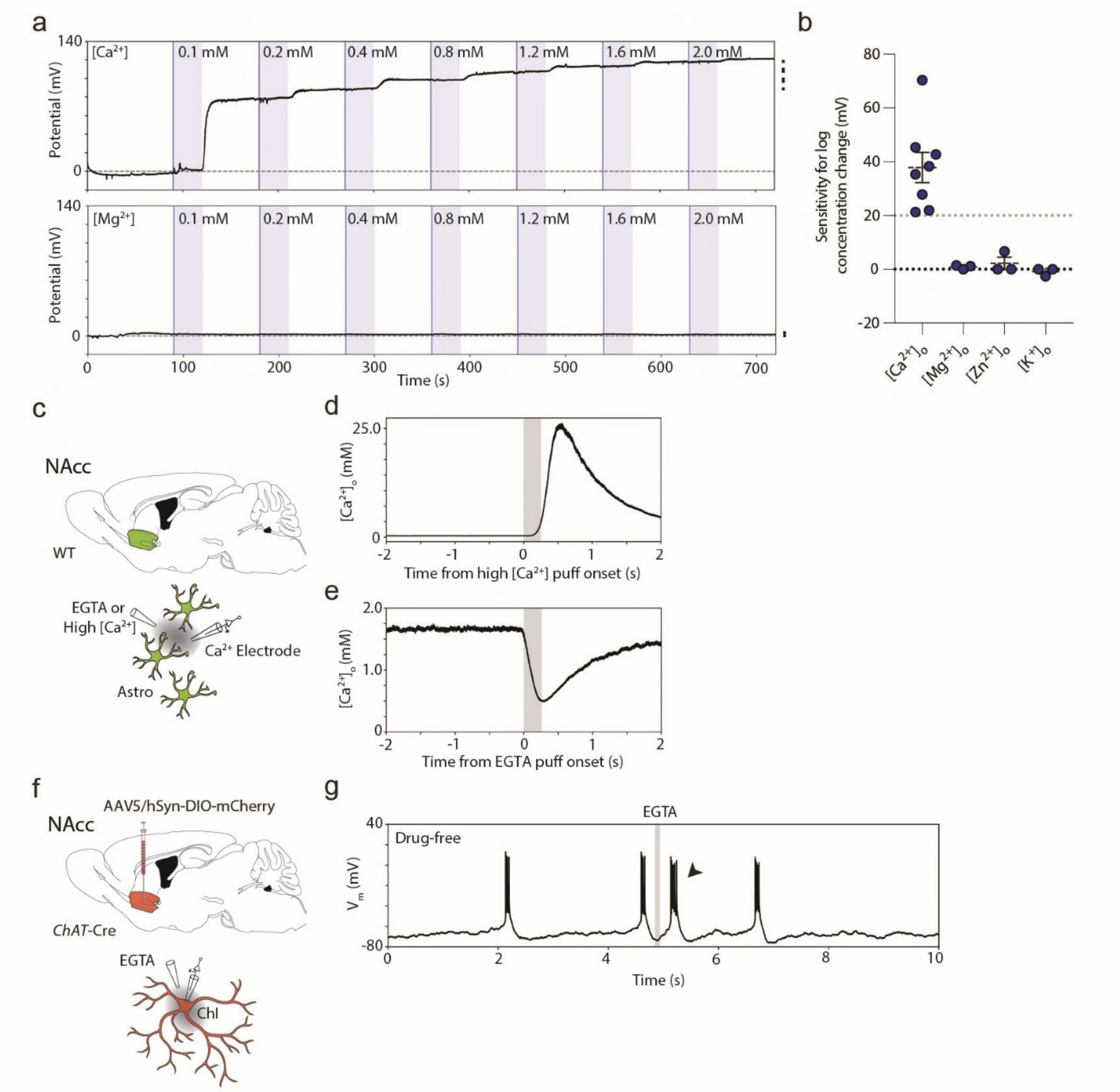


**SUPPLEMENTARY FIGURE S13. $Ca^{2+}$-selective electrodes can sense rapid [$Ca^{2+}$] changes in striatum *ex vivo*. a)** Calibration curves for example $Ca^{2+}$-selective electrode measured as potential jumps in bath application of aCSF with different concentrations (blue columns) of $Ca^{2+}$ (*top*) or $Mg^{2+}$ (*bottom*) over time. Vertical dashed line indicates electrode sensitivity. **b)** Quantification of electrode sensitivity (potential jump from 0.2 mM to 2.0 mM) in aCSF with varying concentrations of [$Ca^{2+}$] ($n$ = 8), [$Mg^{2+}$], [$Zn^{2+}$], or [$K^{+}$] ($n$ = 3). **c)** Experimental overview. $Ca^{2+}$-selective electrodes were placed in wildtype NAcc with an additional glass pipette filled with EGTA (5 mM) or aCSF with 20 mM [$Ca^{2+}$] at ~20 µm distance. **d)** Measured $[Ca^{2+}]_o$ changes in wildtype striatum in response to local puff application (grey column) of aCSF with 20 mM [$Ca^{2+}$]. **e)** Measured $[Ca^{2+}]_o$ changes in wildtype striatum in response to local puff application (grey column) of the calcium chelator EGTA (5 mM). **f)** Experimental approach. **g)** Membrane voltage for a bursting ChI showing an additional bursting response under somatic application of EGTA (5 mM; grey column).

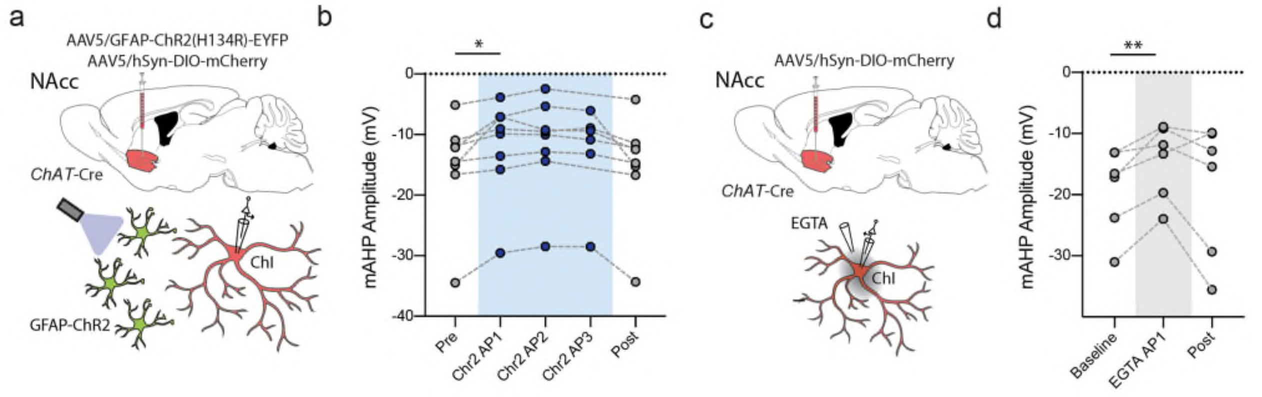


**SUPPLEMENTARY FIGURE S14. Optogenetic activation or extracellular $Ca^{2+}$ chelation diminishes mAHP. a,b)** *ChAT*-Cre mice received intracranial viral expression of hGFAP-ChR2-EYFP and/or hSyn-DIO-mCherry in NAcc, for targeted whole-cell patch clamp recordings of ChIs in NAcc ~3-5 weeks later, as in Figs 3e and 5g. **b,d**) Preliminary analyses suggest post-spike mAHPs in excited ChIs show diminished amplitudes after astrocyte optogenetic activation (from Fig 3e) (b) or local application of a calcium chelator (from Fig 5g) (d), implicating diminished SK-channel function. Amplitude of mAHP was calculated as post-spike mV relative to AP threshold voltage. ** $p < 0.01$, * $p < 0.05$; Paired *t*-tests in (b), (d).

**SUPPLEMENTARY TABLE S1.** Electrophysiological properties of ChIs (*n* = 43(13)).

| **Property** | **mean ± s.e.m.** |
|---|---|
| Input resistance (MΩ) | 184.8 ± 11.1 |
| Resting membrane potential (mV) | -56.3 ± 0.9 |
| Action potential threshold (mV) | -41.3 ± 1.0 |
| Action potential amplitude (mV) | 65.5 ± 1.7 |
| Fast afterhyperpolarization amplitude (mV) | -13.5 ± 0.8 |
| Action potential half-width (ms) | 1.7 ± 0.1 |

**SUPPLEMENTARY TABLE S2.** Electrophysiological properties of pSPNs (*n* = 25(8))

| **Property** | **mean ± s.e.m.** |
|---|---|
| Input resistance (MΩ) | 97.3 ± 12.7 |
| Resting membrane potential (mV) | -80.1 ± 1.5 |
| Action potential threshold (mV) | -35.5 ± 1.7 |
| Action potential amplitude (mV) | 70.1 ± 1.9 |
| Fast afterhyperpolarization amplitude (mV) | -10.8 ± 0.7 |
| Action potential half-width (ms) | 1.3 ± 0.1 |

**SUPPLEMENTARY TABLE S3.** Demographics for human post-mortem brain tissue

| **Patient** | **Age (years)** | **Sex** | **Post-mortem interval (h)** | **Brain Weight (g)** | **Braak Stage** |
|---|---|---|---|---|---|
| HC01 | 82.6 | Male | 45 | 1400 | 2 |
| HC02 | 78.3 | Male | 93 | 1495 | 2 |
| HC03 | 70.2 | Male | 43 | 1540 | 2 |

### Mathematical modelling the proximity of cell types

#### 1. Background

**1.1** We modelled mean minimum centre-centre distances between cholinergic interneurons (referred to here as cell type A) and their nearest neighbour astrocyte (cell type B), and between spiny project neurons (cell type C) and their nearest neighbour astrocyte (B). We used cell densities of $\rho_A$, $\rho_B$ and $\rho_C$ $\mu m^{-3}$ respectively. Table 1 gives, for each cell type the reciprocal of the density, *i.e.* the average striatal volume in which a single cell is found, its cube root $s_X$ which gives the corresponding scale for the distances between cells of type $X$, and the mean of the observed radii for cells of that type.

| Cell type: | A (ChI) | B (astrocyte) | C (SPN) |
|---|---|---|---|
| $1/\rho_X$ | 312500 $\mu m^3$ | 37651 $\mu m^3$ | 6250 $\mu m^3$ |
| $s_X = (1/\rho_X)^{1/3}$ | 67.9 $\mu m$ | 33.5 $\mu m$ | 18.4 $\mu m$ |
| Mean cell radius $r_X$ | 12.5 $\mu m$ | 6.25 $\mu m$ | 6.25 $\mu m$ |

**Table 1: Cell types, densities, and average cell radii**

We modelled numerically the average minimum distance $m(X,Y)$ from a cell of type $X$ to a cell of type $Y$, *i.e.* the distance between a cell of type $X$ to its nearest neighbouring cell of type $Y$, using multiple models.

**1.2** We considered simple models which take as input the densities of two cell-types $X$ and $Y$, and output an estimate of the mean minimum distance from the centre of a cell of type $X$ to a cell of type $Y$. Since the cell diameters are large enough relative to the distance between cells for the assumption that cells are point particles to become an over-simplification, we also consider a variant of the second model which attempts (at least crudely) to take cell diameters into account.

We assume that cell density is homogeneous so that each cell type is distributed uniformly according to their densities. Note that if we were to model this by taking all cells to be uniformly randomly placed, then in any given sample the outcome can result in random samples which are far from homogeneously distributed through the medium. The models we will use will impose a high degree of homogeneity, for simplicity.

#### 2. Model 1

**2.1** The first, simpler, model supposes that the both cell types are distributed in a cubical "lattice packing", that is, at the centres of cubes which completely cover the medium. Each cell of type $X$ must then lie at the centre of a cube of volume $1/\rho_X$ $\mu m^3$, and hence of side-length $s_X = (\rho_X)^{-1/3}$ $\mu m$. If we take units so that $s_X = 1$, then we can suppose that the type-$X$ cells are positioned at the centres of cubes whose vertices are at points ($n_1$, $n_2$, $n_3$) where $n_1$, $n_2$ and $n_3$ are integers. For example, there is a cell at (1/2, 1/2, 1/2) which is the centre of the cube whose vertices have coordinates $\{(a,b,c): a,b,c \in \{0,1\}\}$ and in general, the type $Y$ cells are placed at the points (1/2 + $n_1$, 1/2 + $n_2$, 1/2 + $n_2$) where $n_1$, $n_2$ and $n_3$ are integers.

For a real number $x$, let $[x]$ denote the greatest integer less than $x$ (the expression $[x]$ usually read as "floor $x$") so that $[x] = n$ where $n$ is the unique integer satisfying $n \leq x < n + 1$ and let $\{x\} = x - [x]$ denote the "fractional part" of $x$. If $p$ is a point in $R^3$ with coordinates $(x, y, z)$ and we set $n_x = [x], n_y = [y], n_z = [z]$, then $p$ lies in the cube $[n_x, n_x + 1] \times [n_y, n_y + 1] \times [n_z, n_z + 1]$ and thus its nearest type-$X$ cell has coordinate $(1/2 + n_x, 1/2 + n_y, 1/2 + n_z)$. The distance from $p$ to its nearest type-$Y$ cell is therefore

$$d_Y(p) = \sqrt{\left(\frac{1}{2} - [x]\right)^2 + \left(\frac{1}{2} - [y]\right)^2 + \left(\frac{1}{2} - [z]\right)^2}$$

Now if we let $\beta_Y$ be the magnitude of $s_Y$ in our units where $s_X = 1$, that is, $\beta_Y = s_Y / s_X = (\rho_X/\rho_Y)^{1/3}$. Distributing the type-$Y$ cells in a lattice in the same fashion as the type-$X$ cells corresponds to scaling by $\beta_Y$, so that the type-$Y$ cells are located at the points $\beta_Y(1/2 + m_1, 1/2 + m_2, 1/2 + m_3)$ (where $m_1$, $m_2$, $m_3$ are integers) since these points are the centres of cubes whose vertices have coordinates which are integer multiples of $\beta_Y$.

Now if $\beta_Y$ is rational we may write $\beta_Y = a/b$ for some integers $a$ and $b$ (with $b$ positive and $a$ and $b$ having no common factor). The fractional parts of the coordinates of the cells of type $Y$ have the form $\alpha(n) = \left(n + \frac{1}{2}\right)\beta_Y = \left\{\frac{(2n+1)a}{2b}\right\}$. The function $\alpha(n)$ is then periodic with period $b$, taking the values $\left\{\left\{\frac{a}{2b}\right\}, \left\{\frac{3a}{2b}\right\}, \ldots, \left\{\frac{(2b+1)a}{2b}\right\}\right\}$ hence its values are evenly (though discretely) distributed with mean $1/2$, hence provided $b$ is not too small this is reasonably well approximated by a uniform random variable. On the other hand, if $\beta_Y$ is irrational, by a classical result of Hermann Weyl, $\alpha(n)$ is uniformly distributed over the unit interval. Thus, the mean of the distance $d_Y(X)$ of a cell of type $X$ to its nearest type $Y$ cell is thus approximated by

$$d_3 = E[D] = \int_{[0,1]^3} d_X(x, y, z)\ dxdydz$$
$$= \frac{\sqrt{3}}{8} - \frac{\pi}{48} - \frac{\log(2)}{4} + \frac{\log(1 + \sqrt{3})}{2} \approx 0.4803$$

(Here, as in the rest of the paper, log = $\log_e$, the natural logarithm.) We have thus obtained $d_3$ as an approximation to the mean minimum distance $m(X, Y)$ of a cell of type $X$ to a cell of type $Y$ where $d_3$ has units with $s_X = 1$. In other words, if we let $\pi(X, Y)$ be the model-1 estimate of the mean minimum distance then $\pi(X,Y) = d_3.s_Y = 0.4803\rho_Y^{-1/3}\mu$m.

**2.2** Note that the suitability of the constant $d_3$ in model 1 relies on the assumption that the distribution of the cells of type $Y$ is well-approximated by the lattice modelling it, whereas the term $s_Y = \rho^{-1/3}$ is likely to be more robust, in that it only requires the type-$Y$ cells to be homogeneously distributed.

For example, in the case where the type-$Y$ cells are reasonably homogeneously distributed throughout the medium, but do not exhibit the crystalline arrangement of a lattice, then the factor $d_3$ is likely to underestimate $m(X, Y)$. This is because the mean distance of the points in a unit cube to its centre is smaller than their mean minimum distance to any other point in the cube. It such a context, it is likely to be more appropriate to estimate $m(X, Y)$ by $\Delta_3 \rho_Y^{-\frac{1}{3}}$ where $\Delta_3 \approx 0.661$ is the mean distance between a random pair of points in the unit cube.

**2.3** Note also that for a "thin" medium – that is, a medium which is large in only two spatial directions, such as the 50 μm-thin sections used to observe intercellular distances under fluorescence microscopy distances, it is likely that one will have only a small number of values in the thin spatial ($z$) direction and thus $d_3$ is likely to be inaccurate even where the $Y$ -cells are arranged in a lattice-like manner. At a first estimate, it seems likely that the $x$-$y$ directions should behave according to the 2-dimensional analogue $\Delta_2 \approx 0.5214$ of the above, while in the "thin" $z$-direction, only a few multiples of $\beta_X$ would be observed, and the assumption that they should be well-approximated by a uniform random variable is likely to be questionable.

More generally, it might thus reasonably wish to consider variants of model 1 with a parameter $\lambda$, say

$$\pi_\lambda(X, Y) = \lambda \rho_Y^{-\frac{1}{3}}$$

Of course, the ratio of any two distances predicted by $\pi_\lambda$ is independent of $\lambda$ since

$\pi_\lambda(X_1, Y_2)/\pi_\lambda(X_2, Y_2) = (\rho_{Y2}/\rho_{Y1})^{1/3}$

**2.4** This model also explains why we should not expect $m(X, Y)$ to be symmetric with respect to $X$ and $Y$ : if we assume that the cells of a given type are distributed reasonably homogeneously in the medium, then $m(X,Y)$ should be approximately the same as the distance of an arbitrary point in the medium to its nearest type $Y$ cell. Indeed, the argument above shows that this estimate should be reasonably accurate unless one of $s_X$, $s_Y$ is a small integer multiple of the other.

#### 3. Model 2

**3.1** The second model allows for more variation in the locations of the cells while still requiring them to be reasonably homogeneously distributed. We assume, to begin with, that cell type $X$ is less dense that cell type $Y$, so that $\rho_Y/\rho_X = n_Y > 1$.

The model considers a sample cube with volume equal to $1/\rho_X$ where $\rho_X$ is the density of cell type $X$ and assumes that such a cube will contain one cell of type $X$, and $N_Y$ cells of type $Y$ where $N_Y$ is a Bernoulli random variable which takes the values in $\{n_{Y,}[n_Y] + 1\}$ and has mean $n_Y$. The $N_Y$ type-Y cells and the type-X cell are then modelled by point particles uniformly randomly positioned in the cube, and we estimated the expected minimum distance from the type-$Y$ cells to the type-$X$ cell using a Monte Carlo simulation (see **Appendix,** code available on request**)**.

When $\rho_X > \rho_Y$ we model the medium using a cube of volume $1/\rho_Y$ and calculate the mean of the distances of $N_X$ cells of type $X$ to the cell of type $Y$. Thus model 2 will in this case in effect approximate the average distance between two random points in a cube of side length $s_Y$ . The average distance between two points in a unit cube is known to be

$$\Delta_3 = \frac{4}{105} + \frac{17}{105}\sqrt{2} - \frac{2}{35}\sqrt{3} + \frac{1}{5}\log(1+\sqrt{2}) + \frac{2}{5}\log(2+\sqrt{3}) - 7\pi \approx 0.661$$

**3.2** Model 2 is of course a simplified idealisation. One way in which it is unrealistic is that it assumes the cells are point particles, while the cells' diameter relative to the distances between them make this questionable in some cases. We therefore also ran a slight variant of model 2 where we enforced a lower bound on the minimum distance between cells which we took arbitrarily to be 80% of the larger of the two mean cell radii. In the computer simulation we therefore simply neglected trials where this lower bound on proximity failed. In a simulation with $10^6$ trials, this resulted, for example, in the simulation to calculate m(A,B), in ~9% of trials being neglected.

#### 4. Model 3

**4.1** In addition to the issue of model 2 potentially allowing cell configurations which are not possible because of sizes of the cells, model 2 also assumes that the nearest type X cell to a type Y cell lies in the cube. If, however, one took a random cube C of volume $\rho_Y$ inside the medium, it is possible that the nearest type Y cell to a type X cell lies outside of our cube. In principle it could lie in any of the 26 cubes obtained by translating C by $\pm s_X$ in the direction of any of the axes of the medium. Thus model 2 should overestimate the value of m(X,Y) as a result. One can correct for this overestimate by modelling $3^3$ =27 random cubes and calculating the resulting mean minimum distance of all type Y -cells to the type X cell(s) in the central box. This is what is done in model 3. See **Appendix**.

**4.2** Since the likelihood that the nearest type-Y cell lies outside the cube containing a type-X cell is relatively low, the difference in the predicted values of models 2 and 3 are most significant where the relative densities of the two cell-types is largest, that is, when XY is AB or BA. Just as with model 2, we also ran a variant of model 3 where a lower bound of 80% of the larger of the two mean cell radii was imposed. The predictions of the two variants are listed as models 3 and 3b in the table below.

#### 5. The Predictions

The mean minimum distances, m(X,Y), from a given cell type X to its nearest neighbour Y, as predicted by the models are given in **Table 2**, where we have included the predictions of model 1 using the different parameters $d_3$ and $\Delta_3$. The values in the "*b*" rows for models 2 and 3 are estimates obtained by Monte Carlo simulation with the variants which account for cell diameters using at least $10^5$ trials.

| Model | m(A,B) | m(B,A) | m(C,B) | m(B,C) |
|---|---|---|---|---|
| **1: $\lambda = d_3$** | **16.1 $\mu m$** | 32.6 $\mu m$ | **16.1 $\mu m$** | 8.85 $\mu m$ |
| **1: $\lambda = \Delta_3$** | **22.0 $\mu m$** | 45.0 $\mu m$ | **22.0 $\mu m$** | 12.2 $\mu m$ |
| **2** | **21.7 $\mu m$** | 44.9 $\mu m$ | **22.2 $\mu m$** | 12.0 $\mu m$ |
| **2b** | **23.1 $\mu m$** | 45.3 $\mu m$ | **22.4 $\mu m$** | 12.6 $\mu m$ |
| **3** | **18.3 $\mu m$** | 35.3 $\mu m$ | **17.4 $\mu m$** | 10.0 $\mu m$ |
| **3b** | **19.7 $\mu m$** | 35.7 $\mu m$ | **17.6 $\mu m$** | 10.6 $\mu m$ |

**Table 2: The predicted mean minimum centre-centre distances from each model.**
**Key:** A, ChIs; B, astrocytes; C, SPNs

#### APPENDIX: Details for simulations in Models 2 and 3.

**Model 2.** For model 2 we used computer simulation taking $10^6$ samples to obtain a numerical approximation of the model's prediction for the mean minimum distances $m(X,Y)$. For simplicity in this section, we use units such that the density of cell-type X is equal to 1.

We model the mean minimum distance of a cell of type $X$ to one of type $Y$ where $\rho_Y > 1$ as follows: Let $N_Y$ be a Bernoulli random variable which takes the values $[\rho_Y]$ and $[\rho_Y] + 1$ with probabilities $1 - p$ and $p$ where $p = \{\rho_Y\}$ so that $E[N_Y] = \rho_Y$. We then take $X, Y_1, Y_2, .., Y_{N_Y}$ as independent random variables taking values in $[0,1]^3$, so that each of $Y = (Y_1, Y_2, Y_3)$ $Y_i = (Y_{i,1}, Y_{i,2}, Y_{i,3}) \epsilon [0, 1]^3$ is uniformly distributed.

We then set $D = \min\left\{\sqrt{\sum_{\{k=1}^{3} \left(X_k - Y_{i,k}\right)^2} : i = 1,2, \dots, N_Y\right\}$

and its expectation E[$D$] as our estimate of the mean minimum distance from a cell of type $X$ to a cell of type $Y$. To approximate E[D], the expected value of $D$, we used the random number generator in the NumPy package for Python to produce samples of the random variables $N_Y$, $X_i$, $Y$ and hence to calculate the corresponding value of $D$ and took the resulting average value over $10^6$ simulations.

**Model 3.** Model 3 uses the same paradigm, but generates variables, $N_{Y,1}, \dots, N_{Y,27}$ and, correspondingly, for each $n_{Y,j}$, a set $\left\{Y_k^j : 1 \le j \le 27,\ 1 \le k \le N_{Y,j}\right\}$ of random variables in $(a_1, a_2, a_3) + [0,1]^3$ where $(a_1, a_2, a_3) \in \{-1,0,1\}^3$is given by $j \in \{1,2, \dots ,27\}$ . Then we set, for X, as before, a uniform random variable taking values in $[0,1]^3$,

$$D = \min\left\{\sqrt{\sum_{k=1}^{3}\left(X_k - Y_{i,k}^j\right)^2} : 1 \le j \le 27,\ 1 \le i \le N_{Y,j}\right\}$$

and take $E[D]$ to model the mean minimum distance $m(X,Y)$ from a type-X cell to a type-Y cell. Using the Central Limit Theorem, if, for a sample of size $N$ we write $m_s$ for the sample mean and $\sigma_s$ for the estimator of the standard deviation given by the sample, then provided N is large enough that the Central Limit Theorem ensures E[D] is close to normally distributed, at a 99% confidence level, the expected value $E[D]$ lies in

$$\left[m_s - \frac{3\sigma_s}{\sqrt{N}}, m_s + \frac{3\sigma_s}{\sqrt{N}}\right]$$

When $\rho_Y < 1$ we let $N_X$ be a Bernoulli random variable taking the values $[\rho_Y^{-1}]$ and $[\rho_Y^{-1}] + 1$ with probabilities 1-$p$ and $p = \{\rho_X^{-1}\}$ so that $E[N_X] = \rho_Y^{-1}$. We then generate 27 random variables $\left\{Y_j : 1 \le j \le 27\right\}$ where $Y_j$ takes values in $(a_1, a_2, a_3) + [0,1]^3$ with $(a_1, a_2, a_3) \in \{-1,0,1\}^3$ determined by $j \in \{1,2, \dots ,27\}$, and $N_X$ random variables $\{X_i : 1 \le i \le N_X\}$ each taking values in$[0,1]^3$.

Finally, we set $D_i = \min\left\{\sqrt{\sum_{k=1}^{3}\left(X_{i,k} - Y_{j,k}\right)^2} : j \in \{1,2, \dots ,27\}\right\},\ i \in \{1,2, \dots, N_X\}$

so that each sample in the case where $\rho_Y < 1$ produces $N_X$ trials for $m(X,Y)$ all of which are then averaged over the N samples generated by computer simulation.
